# Conformational and Environmental Determinants of RNA Solvation Dynamics: Roles of Intrinsic Flexibility, Allostery, and Protein Binding

**DOI:** 10.64898/2025.12.24.696429

**Authors:** Amrita Chakraborty, Susmita Roy

## Abstract

Solvation dynamics play a central role in shaping nucleic acid structure, flexibility, and recognition, yet their molecular origins remain poorly understood for RNA, whose diverse architectures and intrinsic conformational plasticity far exceed those of DNA. Here, we present the atomistic, microsecond-scale computational dissection of solvation dynamics in two structurally homologous but functionally distinct viral RNAs—BIV TAR and HIV-2 TAR—in both apo and peptide-bound states mimicking the salt concentration of experimental time-resolved fluorescent spectroscopic measurement. By combining high-temporal-resolution solvation time correlation functions with detailed energy decomposition analyses, we uncover how water, ions, and RNA motions cooperatively shape relaxation across ultrafast to nanosecond timescales. Our results reveal that, unlike DNA, where slow components primarily reflect long-lived hydration and ion condensation, RNA can generate slow solvation decay either through long-lived hydration or through its own internal conformational fluctuations, such as involving spontaneous base-flipping events. Peptide binding modulates this conformational landscape in strikingly system-specific ways: BIV TAR RNA undergoes classical fluctuation quenching, where TAT binding suppresses RNA motions and shifts relaxation toward solvent–RNA compensation, whereas HIV-2 TAR RNA exhibits a non-classical redistribution of solvent–ion–peptide correlations stemming from its weaker and more dynamic binding interface with TAT. The dominant slow decay in HIV-2 apo TAR maps directly onto an allosteric communication channel previously identified from structural analyses, demonstrating that solvation responses can sensitively report on RNA allostery. Together, this study bridges the experimental observations of time resolved fluorescence spectroscopy with mechanistic molecular insight, establishes solvation dynamics as a powerful probe of RNA conformational energetics, and highlights how subtle differences in RNA–protein recognition can imprint distinct signatures on hydration and ion reorganization.

## 1. Introduction

In classical structure–function paradigm, biomolecules such as DNA, RNA, proteins, lipids were considered to traditionally operate within an inert background of water and ions. However, extensive studies over the past decades have revised this notion, demonstrating that water and ions are far from just a passive medium—they dynamically interact with biomolecules, shaping their conformational fluctuations and functional outcomes^1–4^. In proteins and DNA, these coupled dynamics are now known to influence a host of fundamental biological processes^5–9^, ranging from folding and catalysis to molecular recognition and repair. However, molecular-level interaction-based understanding of the explicit dynamics of media relies on underpinning temporal evolution of the solvent’s interaction with a biomolecular solute, and in this regard, Solvation Dynamics^10,11,12^ offers an essential avenue, describing the time-dependent reorganization of the solvent and ionic environment in response to a perturbation of the solute. Upon conformational rearrangement or electronic excitation, the surrounding solvent shell is driven into a nonequilibrium state, prompting ultrafast dipolar reorientation, hydrogen bond network fluctuations, and ion redistribution that relax the system toward a new equilibrium. Experimentally, these processes are captured through time-resolved Stokes shift measurements, such as time-resolved emission spectra^13^ and time-correlated single-photon counting^14^, which monitor the dynamic red-shift in probe emission as the solvent relaxes. Theoretically, Linear Response formalism^15–17^ links these nonequilibrium observables to equilibrium time correlation functions of fluctuation in solute–solvent interaction energy, allowing decomposition into distinct self- and cross-contributions from water, ions, and the biomolecule itself.

For DNA, a wealth of both experimental and computational studies has established multiscale nature of hydration processes for nucleic acid structures, ranging from ultrafast inertial solvent relaxation^18,19^ (tens of femtoseconds) to slower structural rearrangements spanning picoseconds to nanoseconds, associated with several base pairing, base stacking interactions, and conformational plasticity, especially in regions like grooves^20–23^. These dispersed, slow components were traced to long-lived water and ion populations tightly associated with DNA grooves—especially minor groove probe sites—whose local electrostatics create pockets of immobilized solvent molecules and counterion clusters^24,25^. Molecular dynamics simulations and theoretical decompositions have validated these experimental observations and allowed dissection of the underlying molecular causes^15,26,27^. Water residence times around groove-bound ligands often show broad distributions, with many water molecules living in a site-specific manner for hundreds of picoseconds to several nanoseconds—sometimes with over a dozen water molecules displaying residence times >1 ns in single-shell probe regions. Counterion localization, specifically sodium, exhibits structured condensed regions near phosphate backbones and minor grooves^28,29^. Theoretical frameworks, including mode coupling theory^21^ and linear response theory^15–17^, have been deployed to parse the total solvation response into self and cross-component contributions from DNA, water, and ions. Analyses show that the slowest dynamical behaviors arise from coupled fluctuations between DNA and nearby water or between different ion populations, where freezing DNA motion in simulations eliminates much of the ultrafast slow-down, attributing slow solvation phenomena linked to biomolecular flexibility^21,30^. These frameworks also explain why ion-crowding in hypersaline cellular conditions does not substantially alter overall collective solvation dynamics: strong anticorrelation between ion and water energy fluctuations yields a compensatory effect, maintaining net-neutral solvation response even as environmental conditions shift^31^. Other seminal works mapped out sequence-dependent effects^32^, demonstrating that GC pairs in minor grooves interact more strongly with water, while AT-rich regions exhibit unique cation association patterns and greater groove flexibility—a finding with implications for transcriptional regulation and protein-DNA recognition^32–36^. All these pioneering observations not only ties together disparate timescales and components but also highlights how ligand and protein binding, structural disorder, and DNA damage can dynamically reshape the hydration landscape, often with direct consequences for molecular recognition and function.

In contrast, RNA solvation dynamics remain far less explored, despite RNA’s greater structural diversity and functional versatility. Unlike DNA, RNA possesses a unique challenge— higher degree of local flexibility and asymmetry in electrostatics and hierarchical hydration patterns^37,38^ introduce new layers of complexity in interpreting solvation dynamics^39^. In this regard, a time-resolved fluorescence spectroscopic study by Goel et. al with temporal resolution in the range of ∼200 ps, exhibited distinct single and multiexponential relaxation behaviour in BIV TAR RNA and its complexes with cognate TAT peptide, respectively^40^, attributing the involvement of diverse dynamic processes spanning a wide range of timescales. However, one of the key open questions is the extent to which slow solvation responses stem from internal structural fluctuations of RNA itself versus the dynamics of the surrounding water and ionic cloud. For the presence of noncanonical motifs like bulges, internal loops, pseudoknots, and junctions, and single-stranded overhangs, debugging individual contributions from water, ions, and suttle structure-based components is very difficult. Moreover, the highly charged phosphate backbone of RNA draws a dense and fluctuating ionic atmosphere, making it even harder to isolate the solvent-originated contribution from that induced by structural rearrangements. Again, available experimental techniques provide an ensemble-averaged picture, often constrained by temporal resolution. Atomistic molecular dynamics (MD) simulations offer a unique complementary approach, providing temporal resolution down to the femtosecond scale as well as extendable up to sub-picosecond to even nanosecond, further helping disentangle contributions of individual components. Yet, detailed analyses remain scarce for RNA, especially in functionally relevant contexts such as solvation dynamics of functional bulge-containing RNAs that are engaged in protein binding.

This study, to our knowledge is the first exploration of RNA solvation dynamics, computationally leveraging long-timescale (1 μs) atomistic simulations of two viral RNA having similar stem–bulge–loop architecture-BIV TAR and HIV TAR. The solvation dynamics measurement will be compared in both apo and protein-bound states, where cognate protein is an intrinsically disordered 17-mer TAT. By computing solvation time correlation functions with high temporal resolution, we assess the microscopic origins of relaxation components across timescales and dissect the interplay between RNA conformational fluctuations and environmental reorganizations. Comparing two RNAs of similar structural topology but distinct sequence and binding characteristics allows us to highlight how peptide binding modulates the solvation landscape in system-specific ways. Taken together, our study bridges ensemble-averaged experimental measurements with microscopic mechanistic understanding and demonstrates how RNA–protein recognition can be meaningfully interrogated through the lens of solvation dynamics.

## 2. Methods

### 2.1. System Preparation for BIV and HIV-2 and Simulation Protocol

The initial coordinates of the TAR–TAT complexes for BIV (PDB ID: 1BIV ^41^) were obtained directly from the Protein Data Bank, corresponding to the solution NMR structures reported by Patel et al. Among the five available NMR models for the BIV complex, model 1 was selected, as all models exhibit close structural similarity. The corresponding free TAR RNA structure was generated by selectively removing the TAT peptide coordinates from the RNA major groove. An identical procedure was followed for the HIV-2 system, using model 4 of the solution NMR structure (PDB ID: 6MCE ^42^). The BIV PDB entry includes a phosphate cap at the 5′ end of TAR RNA, which was removed prior to simulation.

Both free and bound BIV and HIV systems were subjected to the same simulation workflow. All molecular dynamics simulations were performed using the GROMACS 2018.3 package^43^. System topologies were generated using the AMBER99 force field^44^ with parmbsc0^45^ and χOL3^46^ refinements. Each RNA–protein complex and the corresponding apo RNA were centered in a 12 nm cubic box and solvated with TIP3P ^47^ water. Additional water molecules, along with sodium and chloride ions, were added to ensure charge neutrality and to mimic physiological ionic strength (∼100 mM). After system construction, steric clashes were relieved using steepest-descent energy minimization. Position restraints (force constant: 1000 kJ·mol ¹·nm ²) were then applied to the RNA–protein complexes and RNA-only systems, while ions were kept fixed. A 1 ns NVT equilibration was performed to allow proper ion hydration. The ions were subsequently released and equilibrated for 3 ns under constant volume. The restraints on RNA and protein were gradually reduced (from 1000 → 100 → 10 kJ·mol ¹·nm ²) over an additional 3 ns NVT stage, followed by 500 ps of unrestrained equilibration under NPT conditions. Production molecular dynamics simulations were carried out for 1 µs under the NVT ensemble. (The NVT ensemble was deliberately chosen to maintain bulk salt concentration by preventing box-volume fluctuations.) A leapfrog integrator with a 2 fs timestep was used. Temperature was maintained at 300 K using a Nosé–Hoover thermostat ^48,49^ with a relaxation time of 0.5 ps. During the NPT equilibration stage, pressure was controlled at 1 bar with a Parrinello–Rahman barostat^47^ using a 0.5 ps coupling constant. Periodic boundary conditions were applied throughout. Neighbor lists were updated every 10 steps using a grid scheme. Long-range electrostatics were treated with the Particle Mesh Ewald method ^50^ (Fourier spacing: 0.12 nm, interpolation order: 4). Covalent bond constraints were handled using the LINCS ^51^ algorithm.

### 2.2. Theoretical Framework of Solvation Energy Correlation

Experimentally, this time-dependent reorganization is probed using time-resolved fluorescence Stokes shift (TRFSS) measurements, where the relaxation of solvent polarization around a photoexcited chromophore reports on the local dielectric response. The nonequilibrium solvation response function is then defined as

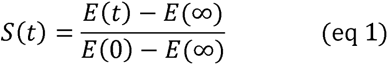

where E(t) is the instantaneous solvation energy following the sudden excitation of the chromophore. Complementarily, in equilibrium simulations, the same dynamics are captured by the energy fluctuation time-correlation function^15–17^

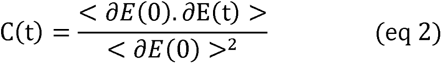

which, under the assumption of linear response, is equivalent to S(t)^15^. Here, ∂E(t) = E(t) - (E), and the brackets ⟨ … ⟩ denote time average over the equilibrium trajectory. Normalization ensures C(0) = 1. Here, E(t) is the instantaneous total interaction energy between the probe and its environment (RNA, water, ions, and protein, if present). The temporal decay of Cs(t) thus reflects the collective relaxation of water molecules, ions, and internal motions of the biomolecule in response to local electrostatic perturbations.

### 2.3. Energy Decomposition Scheme

For a heterogeneous biomolecular environment such as RNA or RNA–protein complexes, as in our case, the total solvation energy fluctuation has been partitioned into contributions from distinct subsystems (e.g., water, counterions, RNA, and protein) following the pioneering work^26,27^ as:

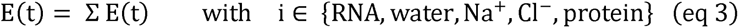

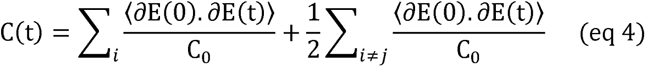

where C_0_ = ⟨ ∂E(0)²⟩ is the total variance. The first term describes self-correlations, while the second captures cross-correlations between subsystems. Positive cross-terms indicate cooperative fluctuations; negative ones correspond to compensatory energy exchanges. Such decomposition^52^ identifies the microscopic origin of slow solvation, usually due to coupling between biomolecular and solvent or ionic motions.

We have further decomposed C(t) to disentangle each component’s (RNA, Water, Ion, RNA and *Protein for complex only*) weightage to total decay function as^27,53^:

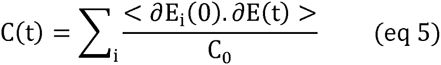

This can guide us which component has the largest variance in fluctuation, contributing the most to the slowness of overall decay function C(t).

### 2.4. Fitting and Characteristic Timescale Estimation

The computed correlation functions C_s_(t) were fitted using a multiexponential decay^10,18^ function:

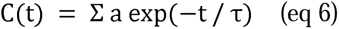

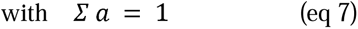

where a and τ denote the amplitude and relaxation time constant of the k-th component. The average solvation timescale expresses how long the overall relaxation persists when multiple exponential components contribute. It can be derived as a weighted sum of the exponential time constants.

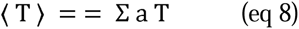

At even longer timescales, a power-law^23^ tail is sometimes observed:

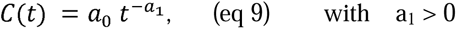

Such behavior implies *scale-free relaxation*, where there is no single dominant timescale. Physically, a power-law decay arises from collective, correlated rearrangements of the extended biomolecular and solvent/ionic environment—such as cooperative ion diffusion, slow domain motion, or large-amplitude hydration-layer fluctuations.

In this work we have employed triexponential function for fitting the data, however, we have shown some of the data fitting exemplifying the power-law behaviour as well.

## 3. Results and Discussions

### 3.1. Peptide-Induced Solvation Dynamics Response in BIV TAR RNA

Early experimental strategies utilize the fluorescent base analog 2-aminopurine (2-AP) at the G11 residue within the bulge region—a location chosen to minimize disruption to local base pairing, allowing for direct interrogation of site-specific solvation dynamics and structural transitions upon ligand interaction. The incorporation of 2-AP provides an exquisitely sensitive probe for capturing changes in solvation environment and associated dynamics triggered by TAT binding near the bulge^40^. To closely mirror these experimental conditions, G11 is selected as the computational Probe site, and the remaining components of the simulation box serve as the surrounding environment in the present study. This enables an atomistic and direct comparison between the simulated solvation dynamics of BIV TAR RNA, both in the ligand-free (apo) state (**Figure 1A**) and in the complexed form bound to the TAT peptide (**Figure 1B**). Computing normalized time-correlation functions Cs(t), which quantify the temporal decay of solvent reorganization around the G11 probe, reveals slower relaxation in the complex state (**Figure 1C)**. By employing Linear Response Decomposition^26,27^, the analysis encompasses all major contributors to the total timescale of solvation dynamics: water, sodium and chloride ions, RNA itself, and, in the bound state, an additional contributor is the peptide, TAT (**Figure 1D-E**). Also, we resolved both self- and cross-correlation terms ^26,27^, parsing the independent and cooperative influences of each component on the observed solvation response (**Figure 2**). Comparing apo and TAT-bound RNA systems reveals distinct solvation signatures: **(i)** In the apo state, rapid relaxation near the probe indicates unobstructed solvent dynamics. Upon TAT binding, interface formation induces a pronounced slowdown, consistent with experiment, manifested by an extended tail in the Cs(t) decay. **(ii)** Dissection of cross-correlation terms reveals enhanced coupling between RNA and solvent in the complex, rather than between solvent and sodium ion, as observed in apo RNA, pinpointing an altered hydration and ion solvation mechanism in the bound state. **(iii)** Consistent with earlier report on DNA solvation dynamics, probe–rest TCF analysis in the case of RNA reveals that water serves as the dominant contributor to the total solvation dynamics for both in its apo and bound state.

**Figure 1:**
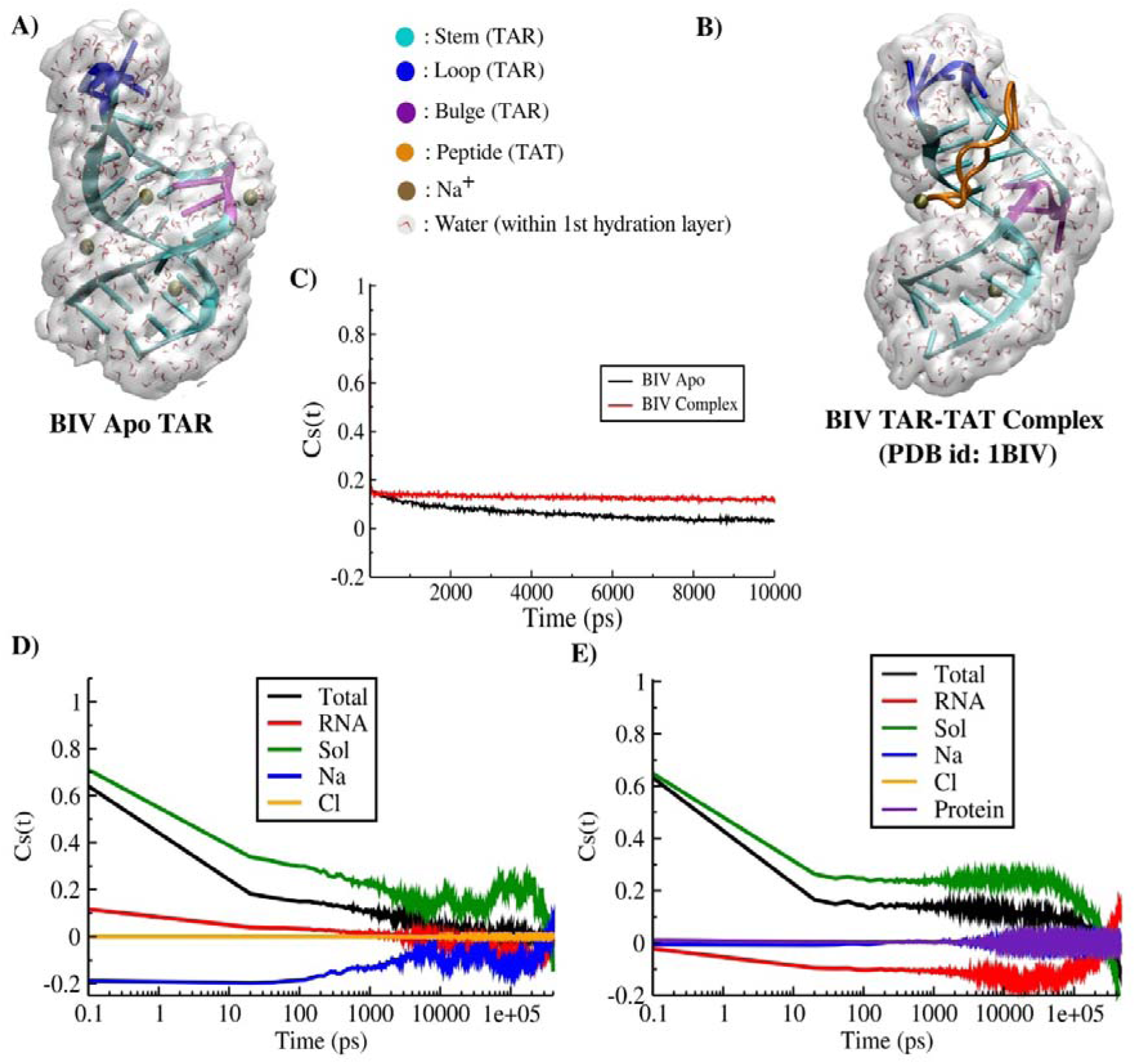
Comparative analysis of solvation dynamics for the BIV TAR RNA in its apo and TAT-bound complex states. (A) and (B) depict the structures of the apo BIV TAR and the TAT-bound complex, respectively, shown as representative snapshots extracted from the corresponding unbiased trajectories. (C) compares the solvation time-correlation functions, Cs(t), for apo and complex systems, represented as decay profiles. Contributions of individual components to the total Cs(t) are shown in **(D)** for the apo state, and in **(E)** for the complex state both presented on semi-logarithmic scale.

**Figure 2:**
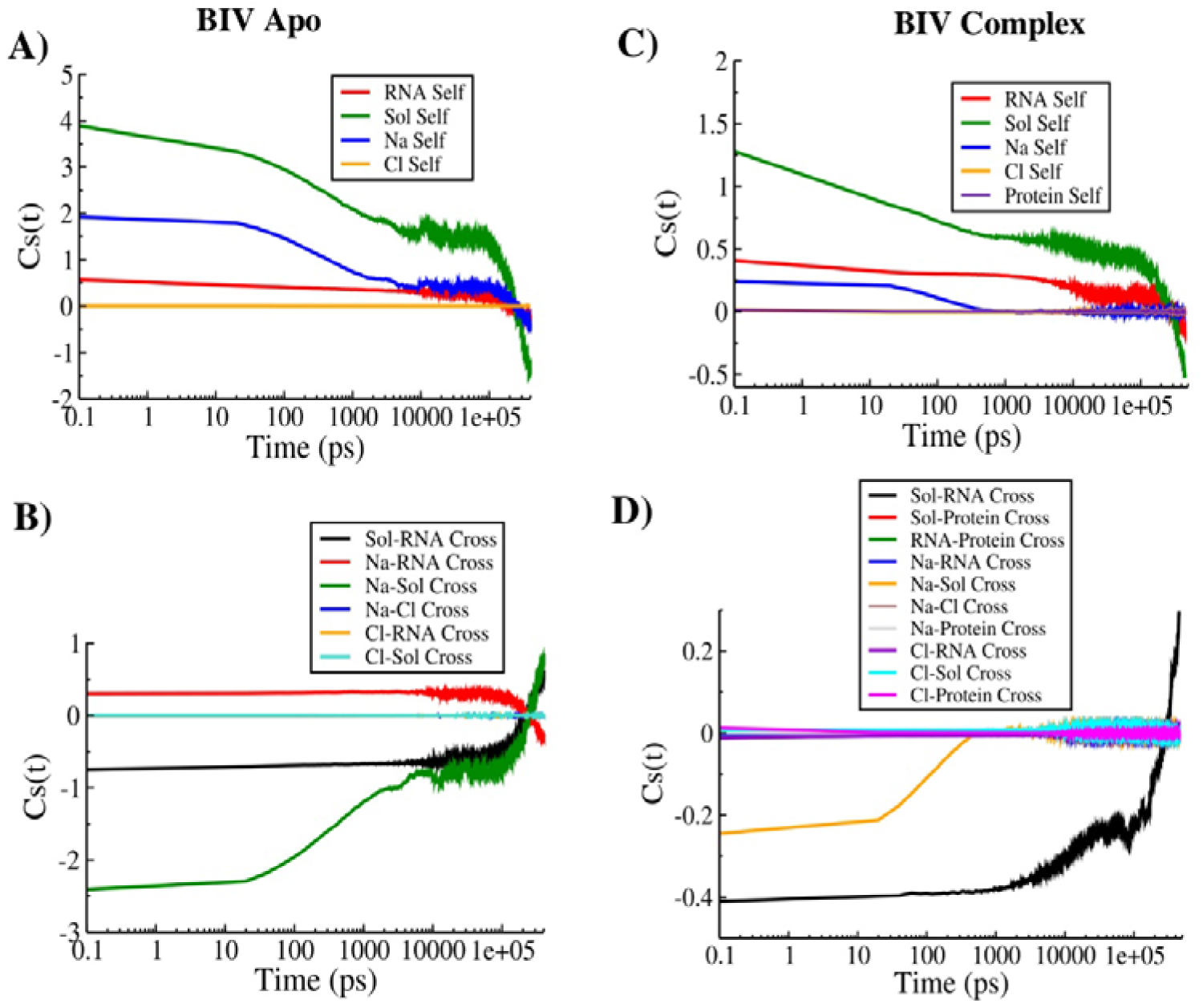
Linear-response decomposition of the BIV TAR RNA decay function into individual self-term and cross-term components. (A) and (B) show the self-terms and cross-terms of the individual components contributing to the decay in the BIV apo TAR. (C) and (D) present the corresponding terms for the TAR–TAT complex.

At this stage, we aimed to understand microscopically how protein association modulates RNA solvation behavior at experimentally relevant probe sites. To this end, we systematically reviewed total (Probe-Rest) energy profiles and energy fluctuations (ΔE), monitoring contributions from RNA, water (solvent), sodium, chloride ions, and protein (**Figure 3**) and it became immediately apparent that TAT binding could not remodel the mean energy but instead alters the fluctuation amplitudes for solvent and ion terms, suggesting a reorganization of local environmental dynamics around the bulge-binding interface. This provides direct comparative insight into the shift from more frequent water–ion anticorrelation in the apo state to less frequent anticorrelation in the complex.

**Figure 3:**
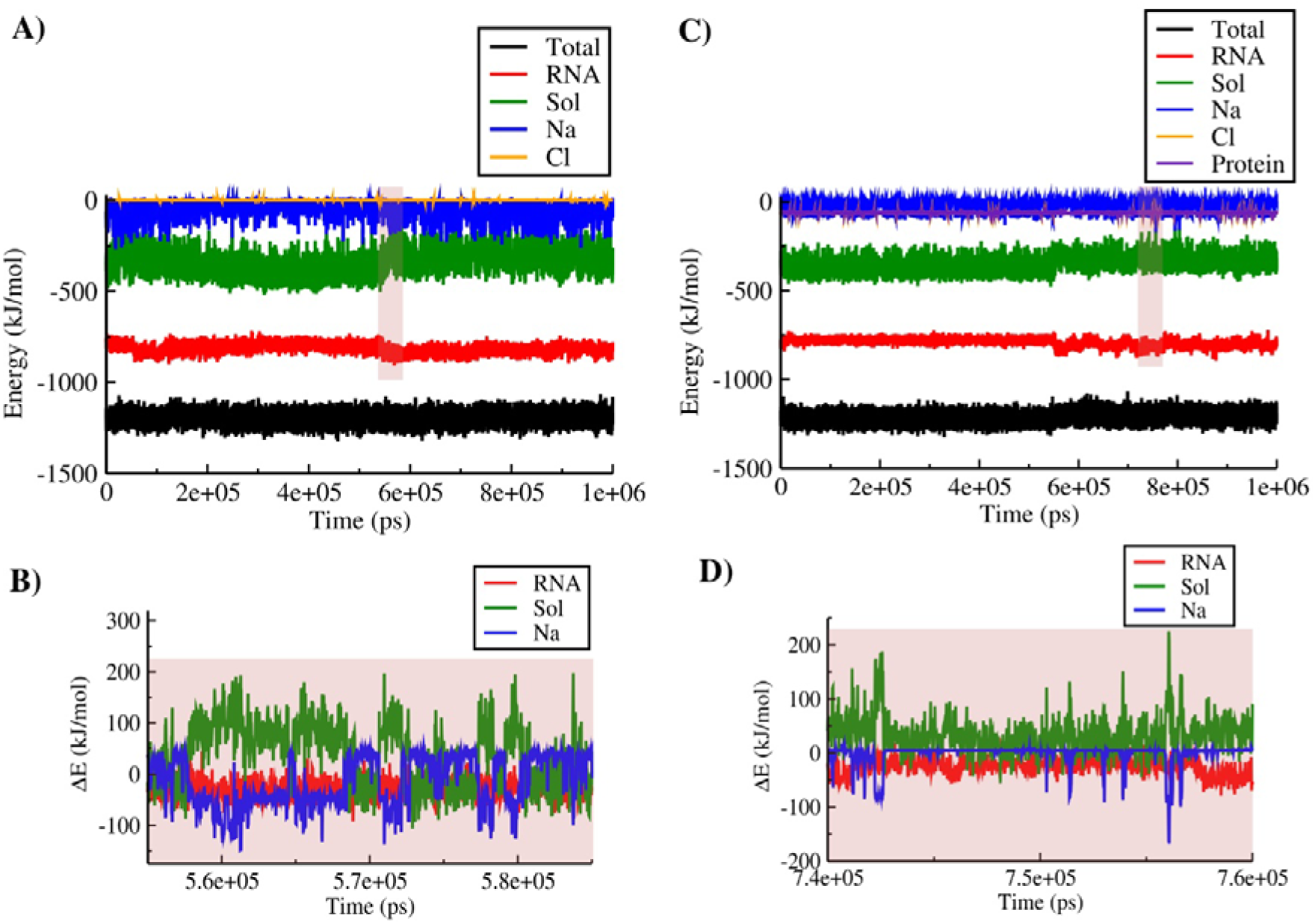
Solvation energy decomposition for the BIV TAR RNA, highlighting contributions from different environmental components in both the apo and complex states. (A) Probe–rest interaction energy for the BIV apo RNA, shown both as the total contribution and as its decomposition into individual components. (B) Magnified view of panel (A), highlighting the fluctuations in interaction energy (ΔE) for the RNA, solvent, and Na^+^ ion components. (C) and (D) Corresponding plots for the BIV TAR–TAT complex, analogous to panels (A) and (B), with an additional contribution arising from the TAT peptide.

How this solvent-ion reorganization pattern induced by TAT binding drives the transition from a dynamically open (apo) to a locally immobilized (complexed) solvation shell requires examining spatial and temporal signatures of water and Na^+^ ions around the BIV TAR RNA, in both its apo and complex form. The radial distribution of water around the RNA reveals a pronounced first-shell peak in the apo RNA, reflecting a well-organized hydration layer. In contrast, upon complex formation with TAT, the first-shell peaks are significantly attenuated, indicating partial dehydration of the bulge/loop interface as the peptide inserts into the major groove. This reduction in local water density represents the classic picture of water release to the bulk during biomolecular association. Upon TAT binding, both water and sodium g(r) profiles flatten and broaden, indicative of disrupted structural order (**Figure 4A-B**). The reduced sodium density suggests that TAT effectively competes for ion-binding hotspots, thereby altering the electrostatic landscape and potentially affecting RNA stability.

**Figure 4:**
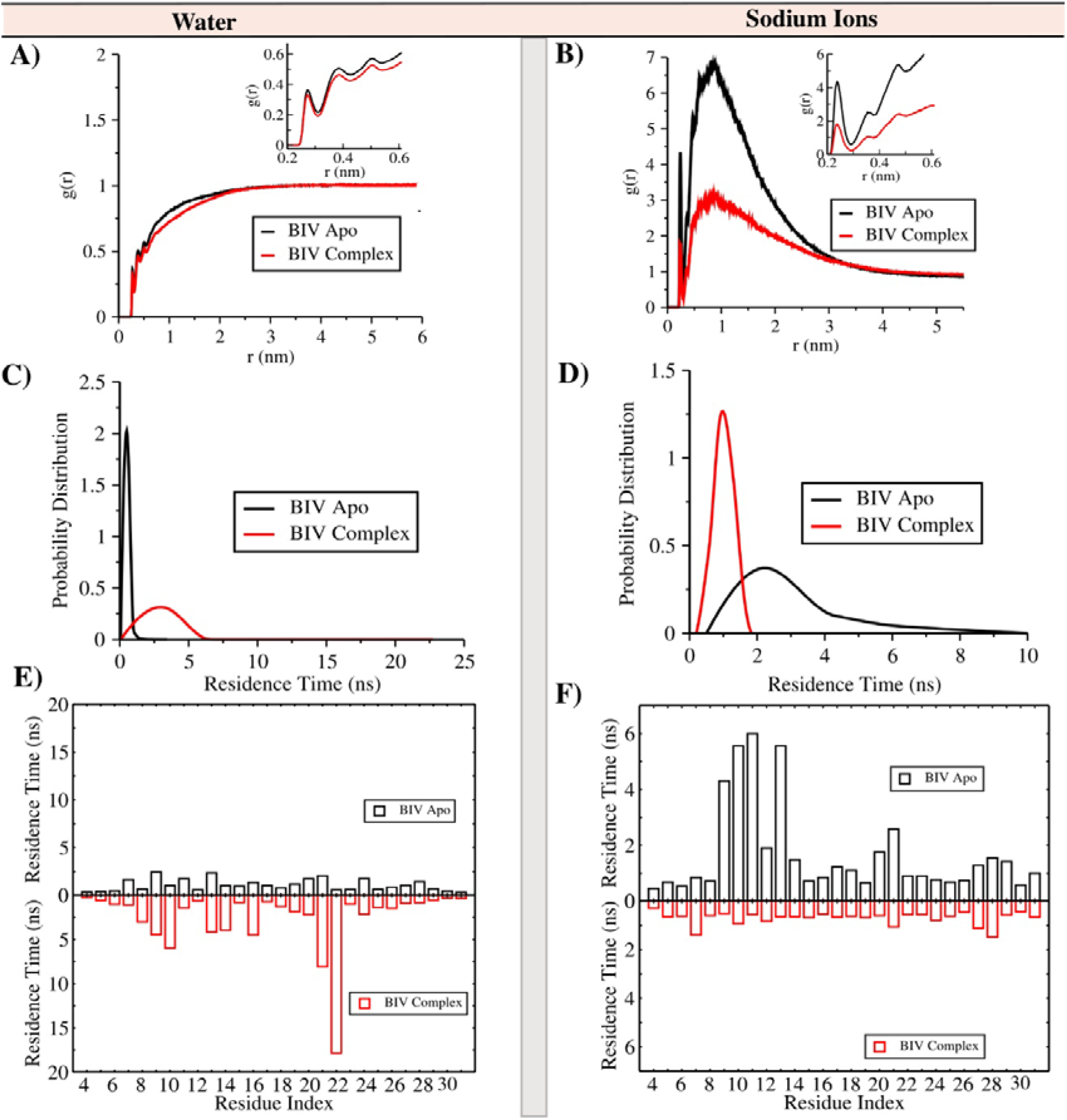
Characterization of the water and ion (Na^+^) environment surrounding the RNA duplex in the apo and TAT-bound states of BIV TAR. (A) and (B) show the radial distribution functions of water and Na^+^ ions, respectively, around all RNA heavy atoms, compared between the apo and complex forms. (C) and (D) present the distributions of residence times for water and Na^+^ ions within the first hydration shell (up to 3.5 Å) of the RNA in both states. (E) and (F) illustrate how the maximum residue-wise residence times of water and Na^+^ ions in the first hydration shell of the apo RNA are altered upon TAT binding.

Kinetic aspects of solvent and ion dynamics are captured by the residence time distributions (**Figure 4C-D**). Water in the apo RNA typically exhibits sub-nanosecond residence times, with only a small fraction persisting beyond 1 ns. Upon peptide binding, however, the tail of the distribution extends to several nanoseconds (**Figure S1**), with a distinct population surviving up to ∼3 ns. This shift underscores the emergence of longer-lived hydration near the protein–RNA interface. In contrast, Na^+^ residence times show an opposite trend: while the apo RNA accommodates ions with multi-nanosecond lifetimes, these long-lived states are diminished in the complex, consistent with peptide competition for the bulge major groove binding site.

Finally, residue-wise decomposition of water (**Figure 4E**) and Na^+^ (**Figure 4F**) residence times highlights the specific RNA regions most affected. For water, nucleotides near the bulge and loop exhibit a modest to pronounced increase in residence lifetimes upon peptide binding, pointing to the stabilization of hydration networks in these flexible domains. For Na^+^, the apo state shows extended binding near the bulge nucleotides, which is substantially weakened in the complex, again consistent with the peptide displacing or reorganizing the local ionic environment. Thus, TAT peptide remodels the local environment of BIV TAR RNA, particularly the bulge in the major groove, the principal TAT binding site central to viral replication and structural specificity. Water becomes more structured and long-lived, while Na^+^ ions are partially excluded upon complex formation.

### 3.2. Timescale Decomposition of Solvation Response in Apo RNA and RNA-Protein Complex

As noted earlier, the 20 ps-resolution trajectories could not fully capture the sharp ultrafast decay component, since ∼70–80% of the correlation was already lost within the first 20 ps in both apo and complex states. However, because our current focus is on understanding the origin of the long-time behavior, we restrict the timescale analysis here to the 20–ps–resolution data (**Figure 5**). Only equilibrated segments of the trajectories were used for all fitting procedures, and the extracted timescales are summarized in **Tables 1–2**. We have fitted the decay plot of TAR-TAT complex using power law fitting equation (**Figure S2, Table S1**) as well, as it is behaving way slower than apo TAR.

**Figure 5:**
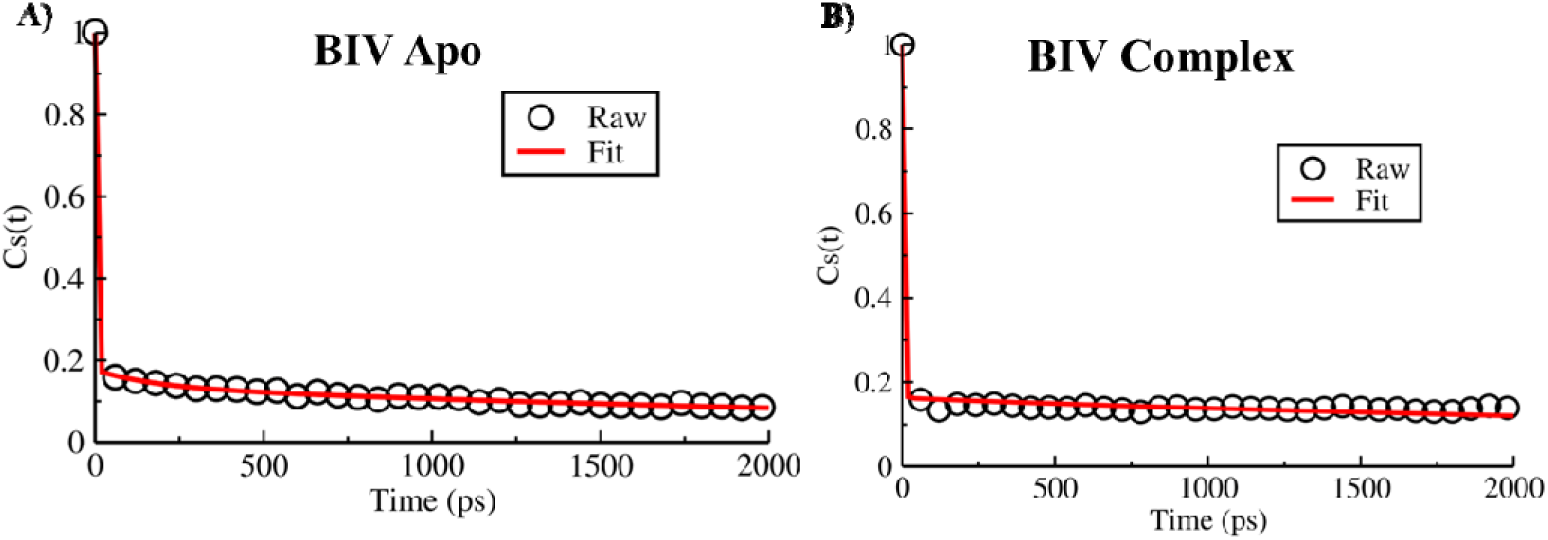
Triexponential fitting of the solvation decay profiles for the apo and complex forms of BIV TAR. (A) Decay profile of the apo TAR RNA, fitted up to 2 ns using a triexponential function. (B) Corresponding decay profile for the TAR–TAT complex, fitted using the same triexponential model.

**Table 1:** Fitting parameter for BIV Apo (Tri-exponential)

| **a1** | **a2** | **a3** | **T1** | **T2** | **T3** | **<T>** |
| --- | --- | --- | --- | --- | --- | --- |
| **0.824** | **0.042** | **0.134** | **0.72 ps** | **178.66 ps** | **4231.36 ps** | **575.10 ps** |

**Table 2:**
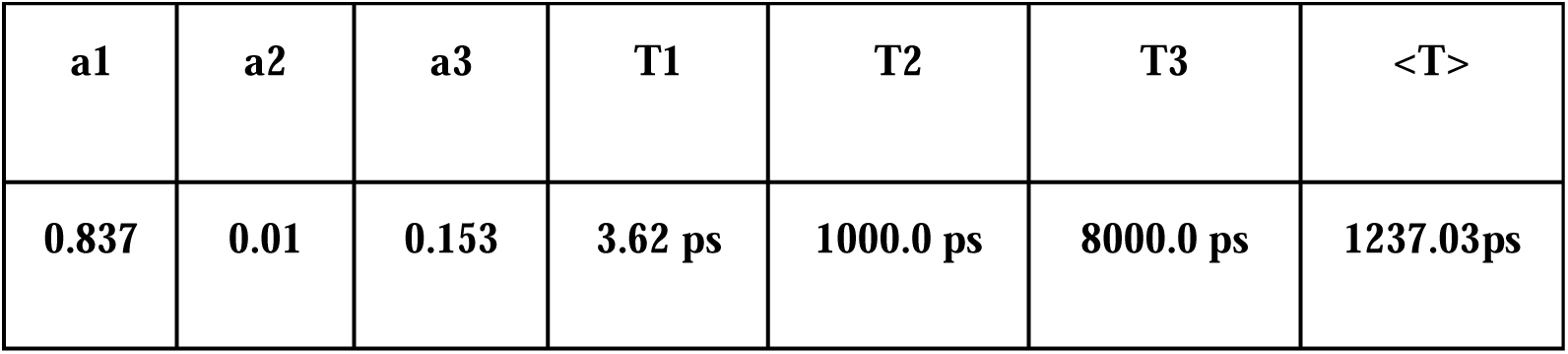
Fitting parameter for BIV TAR-TAT Complex (Tri-exponential)

| **a1** | **a2** | **a3** | **T1** | **T2** | **T3** | **<T>** |
| --- | --- | --- | --- | --- | --- | --- |
| **0.837** | **0.01** | **0.153** | **3.62 ps** | **1000.0 ps** | **8000.0 ps** | **1237.03ps** |

Fitting of the 20 ps-resolution data yields average relaxation times that lie closer to the experimentally observed nanosecond regime, with a multi-exponential hierarchy that reflects contributions from both water–ion reorganization and RNA motions. Importantly, although exact quantitative agreement is not expected—given the restricted ensemble sampling of a single-molecule trajectory and inherent differences between experimental and simulation resolutions—the hierarchy of timescales and the relative slowing upon complex formation are fully consistent with experimental observations. Later on, to overcome the issues of faster uncorrelation of data obtained at 20ps resolution, we carried out additional 20 ns simulations sampled at a finer resolution (1ps), thereby extending the accessible timescales from femtosecond regime (**Figure S3**, **Table S2-S3**).

### 3.3. How RNA’s Conformational Transitions Shape Solvent Relaxation in a System-Specific Manner

Although decomposition of the BIV TAR probe-rest TCF shows that the solvent provides the dominant contribution across all timescales—consistent with its larger variance—an important observation emerges when examining different trajectory segments: the BIV complex exhibits distinct decay patterns (**Figure S4**), presumably driven by underlying structural degrees of freedom. This redirects our attention to the coupling between water dynamics and RNA structure. Notably, the solvent contribution is not purely ultrafast; its self-term decays across the entire ps–µs window, reflecting contributions from both bulk water and groove-confined hydration. RNA self-fluctuations, though smaller in magnitude, remain correlated over long times, generating a persistent background component. Importantly, the solvent–RNA cross-term becomes strongly negative in the complex, indicating anti-correlated fluctuations between RNA breathing motions and compensatory solvent rearrangements. Thus, although water numerically dominates the variance, the overall decay pattern is inseparably shaped by RNA–solvent coupling. Segmenting the trajectory (**Figure 6A**) further reveals that when no local structural changes occur, relaxation remains fast and water-dominated (**Figure 6B**), whereas segments involving a local transition near the probe produce a markedly slower decay, with RNA fluctuations contributing substantially at late times (**Figure 6C**), highlighting how the solvent relaxation is modulated by the RNA’s conformational subspace.

**Figure 6:**
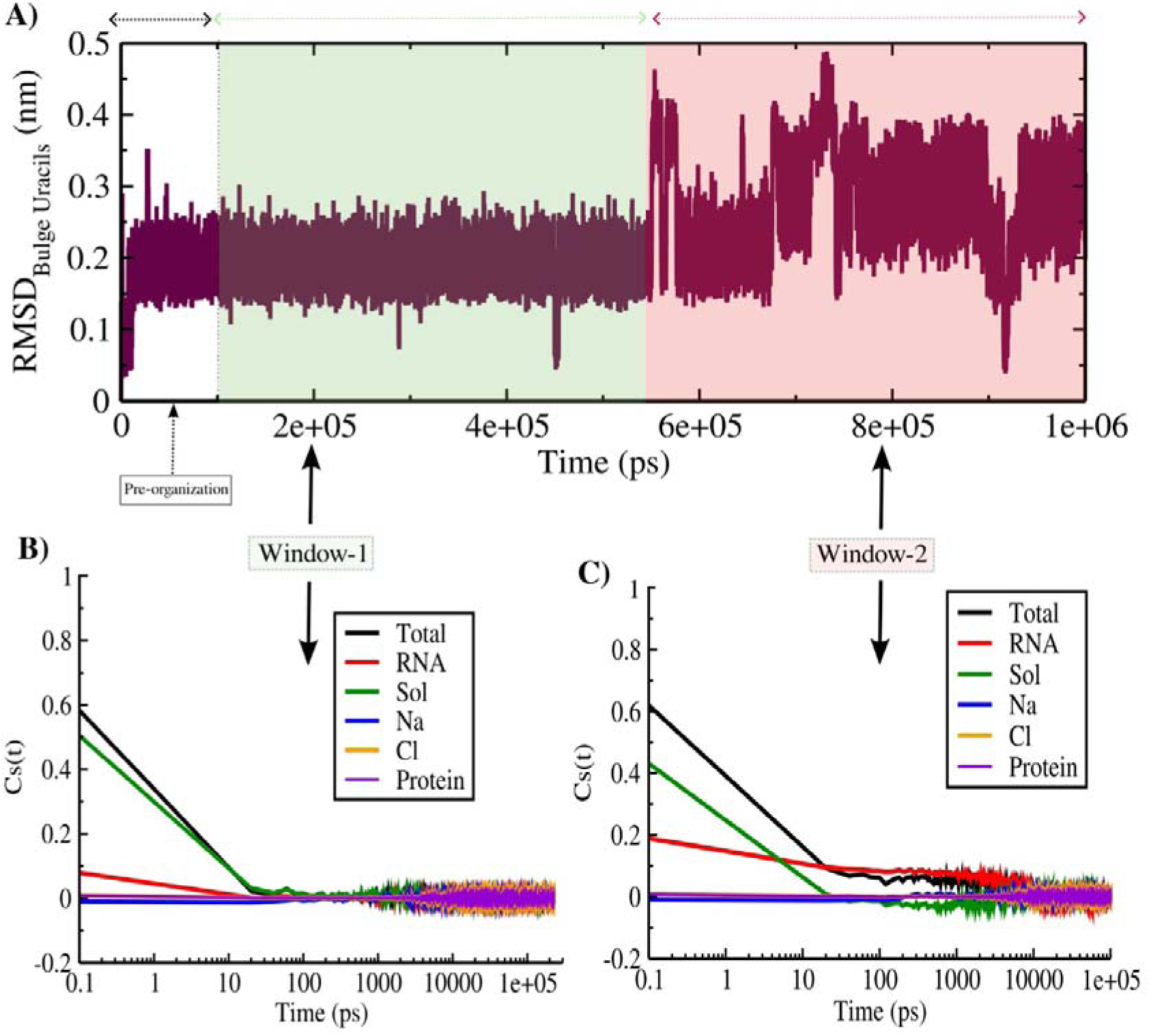
How distinct RNA conformational transitions modulate solvent relaxation. (A) RMSD of the bulge uracils—the residues nearest to the probe—plotted along the trajectory. The green and red shaded regions denote two segments/windows of a single trajectory, segmented based on conformational changes evident from the RMSD analysis. (B) and (C) Window-specific decompositions of the component-wise contributions to the solvation time correlation function for the green and red trajectory windows, respectively. Note the distinct conformational fluctuations between the two windows.

With this understanding, we now turn to the HIV system, which shares an analogous stem-bulge-loop RNA hairpin topology. In the apo state, HIV-2 TAR RNA is known to undergo pronounced groove breathing motions driven by loop–bulge communication, propelled by a specific base flipping from loop (A35). If a seemingly subtle fluctuation in BIV complex can markedly alter the interplay between RNA and water, then the more extensive breathing motions in HIV TAR are expected to leave an even stronger imprint on its solvation dynamics. A central question, therefore, is how these breathing modes are quenched upon TAT engagement, and how this suppression of RNA dynamics reshapes the overall solvation decay in the HIV complex, rendering it distinct from the structurally related BIV counterpart

To probe the solvation dynamics associated with pre- and post-TAT bound form of HIV-2 TAR (**Figure 7A-B**), we strategically select U23 from the bulge as the solvation probe, since it simultaneously reports on loop–bulge stacking driven by A35 flipping and on direct TAT binding, where U23 is a major contact site. Importantly, U23 shows no erratic fluctuations, satisfying the prerequisites for equilibrium response measurements. Parallel to the BIV analysis, we first computed the total Probe-Rest interaction energies and their component-wise contributions in both apo and complex states. The zoomed-in view of the energy fluctuations indicate that the solvent is either correlated or anticorrelated with the biomolecules (RNA/Peptide) or ion (**Figure S5**). The decay profile for apo versus complex reveal that HIV-2 Apo undergoes a slower relaxation (**Figure S5, Table S4, S5**). The comparatively slower solvation dynamics observed for HIV-2 apo (**Figure 7A**) compared to BIV apo can be attributed to the distal-loop base flipping of A35, which drives periodic groove-breathing motions (**Figure 7B-D**), and the corresponding energy fluctuations occur on comparable timescales (**Figure S6**). Component-wise decomposition clearly shows that RNA dominates the long-time regime, confirming that intrinsic conformational fluctuations are directly imprinted onto solvation dynamics (**Figure 7E**).

**Figure 7:**
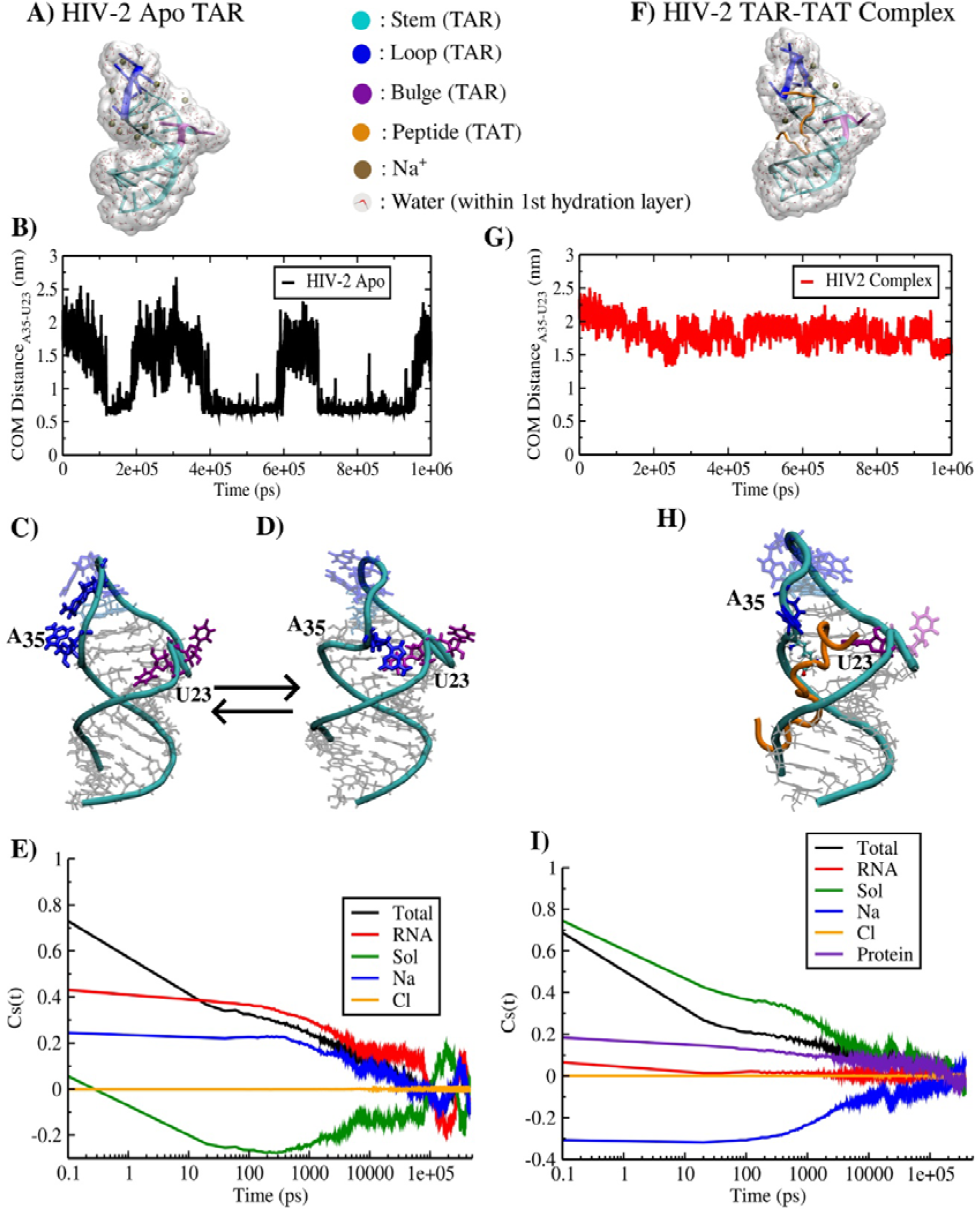
Connection between intrinsic flexibility of RNA and solvation dynamics comparing HIV-2 apo TAR and TAR-TAT complex. (A) Structure of the apo HIV TAR RNA, shown as a representative snapshot taken from an unbiased trajectory. (B) Centre-of-mass (COM) distance between residues A35 (loop) and U23 (bulge) in the apo state. (C) and (D) Depictions of the dynamic “breathing” between two conformational states, driven by reversible base stacking between A35 and U23. (E) Individual component-wise contributions to the total Cs(t) for the apo system. (F) Structure of the TAT-bound HIV TAR RNA as reported by Pham et al. (PDB id: 6MCE^42^). (G) COM distance between residues A35 and U23 in the TAR–TAT complex, corresponding to the structure shown in (H). (I) shows individual contributions to the total Cs(t) in the complex form.

In contrast, the TAT-bound form (**Figure 7F**), which lacks the apo-like dynamic stacking associated with A35 (**Figure 7 G-H**), exhibits the classic picture where solvent provides the dominant contribution (**Figure 7I**). Notably, linear response decomposition of Cs(t) into self-and cross-terms of each component pinpoints subtle yet functionally meaningful distinctions arising from TAT. Whereas the BIV complex exhibits negligible TAT self-contribution, the HIV-2 complex shows a substantial TAT self-term—ranking after solvent and sodium—while the RNA self-term is comparatively suppressed. Among the cross-terms, the solvent–sodium pair dominates, as in BIV, and sodium–TAT emerges as the second largest contributor (**Figure S7**).

## 4. Conclusion

The results of this study return to—and substantially extend—the central question posed in the introduction: to what extent do slow solvation responses in nucleic acids arise from solvation reorganization versus internal conformational fluctuations? While decades of work on DNA established that long-lived hydration, ion residence, and groove-specific dynamics dominate slow components of solvation relaxation^3,54–58^ ,for RNA^59^, it has remained scarce due to its far greater conformational heterogeneity, asymmetric electrostatics, and motif-dependent hydration patterns^39,60–62^.

By dissecting the solvation response of two architecturally similar but mechanistically distinct TAR RNAs—BIV and HIV-2—we show that RNA solvation dynamics do not conform to a single universal framework. Instead, they emerge from a context-dependent interplay between environmental fluctuations and intrinsic RNA motions. In BIV TAR, even a subtle change in conformational fluctuation in the apo state is sufficient to shift the slowest components of solvation from water–Na anticorrelation to a predominantly RNA–solvent compensated regime upon TAT binding. This transition resembles the “classical” behaviour extensively documented for DNA hydration and ligand binding, where biomolecular motions are quenched and the solvent–ion cloud reorganizes cooperatively. Consistent with this paradigm, the BIV complex shows: negligible peptide self-correlation, diminished solvent–Na anticorrelations, and a dominant RNA–solvent compensatory mode. This reflects the tightly locked geometry of the BIV TAR–TAT interface, as also indicated by spectroscopic studies^40^. In HIV-2 TAR, the behaviour is fundamentally different. Its apo RNA exhibits an even slower decay than BIV—driven directly by long-timescale intrinsic motions associated with A35-mediated loop–bulge communication^63^. These internal dynamics leave a strong fingerprint on the environment, generating large RNA self-correlations absent in BIV. Upon TAT binding, these motions are only partially quenched: the weaker-binding peptide retains significant mobility and couples dynamically to both solvent and sodium ions. As a result, the HIV-2 complex exhibits a non-classical solvation hierarchy, where: solvent–Na dominates, Na^+^ –TAT emerges as the second largest contributor, and RNA–solvent anticorrelation drops to fourth—a qualitative inversion of the BIV pattern. This ranking signals a highly dynamic, partially hydrated recognition interface, wherein the probe lies.

Our findings also connect directly to RNA allostery. Earlier structural work^63,64^ identified a prebound-state allosteric channel in HIV-2 TAR involving transient A35–U23 stacking. In this study we detect the same communication pathway that responds through slow environmental relaxation: the apo-state solvation decay is governed by fluctuations tied to this stacking transition. Thus, solvation response functions—traditionally treated as hydration probes—emerge here as sensitive detectors of allosteric propagation^65^ in RNA, in agreement with emerging views of RNA as a mechanically coupled network^66^. Consistent with early observation of prevailing biomolecular dynamics caused by abasic damage dominating solvation dynamics in DNA^67^, for the first time here in this study RNA conformational transition has been attributed to the long timescale of overall solvation response.

Finally, differences in hydration **(Figure S8)** provide a mechanistic basis for the well-known disparity in TAR–TAT affinities^68,69^. HIV-2 retains a larger population of long-residence waters near the groove, whether those screen peptide insertion or compensate for its weaker stabilizing interactions-this question is deemed a separate study. In contrast, BIV integrates high-residence waters, but less in number into its compact interface, enabling tighter and more cooperative recognition—a mechanism consistent with hydration-controlled affinity in nucleic acid–protein complexes^56,70,71^.

In summary, our study reveals that:

i. Slow solvation in RNA can be dominated by intrinsic conformational dynamics, unlike the largely environment-driven behavior in DNA.
ii. Peptide binding reshapes solvation differently depending on the RNA–protein interface, producing either classical quenching (BIV) or non-classical redistribution of environmental correlations (HIV-2).
iii. Solvation response functions serve as powerful, underappreciated reporters of RNA allostery and recognition pathways.

These findings open a new mechanistic framework for interpreting how RNA hydration, flexibility, and protein binding co-regulate molecular recognition at the nanoscale.

## Availability of Data and Materials

All data and codes used in the analysis are available from the corresponding author to any researcher for purposes of reproducing or extending the analysis under a material transfer agreement with IISER-Kolkata, India.

## Author contributions

Conceptualization: S. R. Data curation: A.C. Formal analysis: A.C., S. R. Funding acquisition: S. R. Methodology: S. R., A.C. Project administration: S. R. Resources: S. R., Validation: S. R., A.C. Writing – original draft: S. R., A. C. Writing and editing: S. R., A.C.

## ACKNOWLEDGEMENTS

SR acknowledges support from the Department of Biotechnology (DBT) (Grant No. BT/12/IYBA/2019/12 and BT/PR40192/BTIS/137/69/2023). AC acknowledges DST INSPIRE fellowship.

## References

1 N. Nandi, K. Bhattacharyya and B. Bagchi, Dielectric relaxation and solvation dynamics of water in complex chemical and biological systems, Chem Rev, DOI:10.1021/cr980127v.

2 P. Ball, Water as an active constituent in cell biology, 2008, preprint, DOI: 10.1021/cr068037a.

3 D. Laage, T. Elsaesser and J. T. Hynes, Water Dynamics in the Hydration Shells of Biomolecules, Chem Rev, DOI:10.1021/acs.chemrev.6b00765.

4 S. Khodadadi, J. H. Roh, A. Kisliuk, E. Mamontov, M. Tyagi, S. A. Woodson, R. M. Briber and A. P. Sokolov, Dynamics of biological macromolecules: not a simple slaving by hydration water, Biophys J, DOI:10.1016/j.bpj.2009.12.4284.

5 B. Bagchi, Water dynamics in the hydration layer around proteins and micelles, 2005, preprint, DOI: 10.1021/cr020661+.

6 S. K. Pal, J. Peon, B. Bagchi and A. H. Zewail, Biological water: Femtosecond dynamics of macromolecular hydration, Journal of Physical Chemistry B, DOI:10.1021/jp0213506.

7 B. Jana, S. Pal, P. K. Maiti, S. T. Lin, J. T. Hynes and B. Bagchi, Entropy of water in the hydration layer of major and minor grooves of DNA, Journal of Physical Chemistry B, DOI:10.1021/jp061588k.

8 S. Roy and B. Bagchi, Solvation dynamics of tryptophan in water-dimethyl sulfoxide binary mixture: In search of molecular origin of composition dependent multiple anomalies, Journal of Chemical Physics, DOI:10.1063/1.4813417.

9 S. Wang, J. Gao and X. Chu, Residue-Specific Structural and Dynamical Coupling of Protein and Hydration Water Revealed by Molecular Dynamics Simulations, Biomolecules, 2025, 15, 660.

10 B. Bagchi and B. Jana, Solvation dynamics in dipolar liquids, Chem Soc Rev, DOI:10.1039/b902048a.

11 D. Roy and M. Maroncelli, Simulations of solvation and solvation dynamics in an idealized ionic liquid model, Journal of Physical Chemistry B, DOI:10.1021/jp301359w.

12 A. Chandra and B. Bagchi, Molecular theory of solvation and solvation dynamics of a classical ion in a dipolar liquid, Journal of Physical Chemistry, DOI:10.1021/j100356a023.

13 D. P. Millar, Time resolved fluorescence spectroscopy, Curr Opin Struct Biol, DOI:10.1016/S0959-440X(96)80030-3.

14 W. Becker, A. Bergmann, M. A. Hink, K. König, K. Benndorf and C. Biskup, Fluorescence Lifetime Imaging by Time-Correlated Single-Photon Counting, Microsc Res Tech, DOI:10.1002/jemt.10421.

15 M. Maroncelli and G. R. Fleming, Computer simulation of the dynamics of aqueous solvation, J Chem Phys, DOI:10.1063/1.455649.

16 B. B. Laird and W. H. Thompson, On the connection between Gaussian statistics and excited-state linear response for time-dependent fluorescence, Journal of Chemical Physics, DOI:10.1063/1.2747237.

17 E. A. Carter and J. T. Hynes, Solvation dynamics for an ion pair in a polar solvent: Time-dependent fluorescence and photochemical charge transfer, J Chem Phys, DOI:10.1063/1.460431.

18 R. Jimenez, G. R. Fleming, P. V. Kumar and M. Maroncelli, Femtosecond solvation dynamics of water, Nature, DOI:10.1038/369471a0.

19 S. K. Pal, L. Zhao and A. H. Zewail, Water at DNA surfaces: Ultrafast dynamics in minor groove recognition, Proc Natl Acad Sci U S A, DOI:10.1073/pnas.1433066100.

20 N. Pal, H. Shweta, M. K. Singh, S. D. Verma and S. Sen, Power-law solvation dynamics in G-quadruplex DNA: Role of hydration dynamics on ligand solvation inside DNA, Journal of Physical Chemistry Letters, DOI:10.1021/acs.jpclett.5b00653.

21 B. Bagchi, Anomalous power law decay in solvation dynamics of DNA: A mode coupling theory analysis of ion contribution, Mol Phys, DOI:10.1080/00268976.2014.904943.

22 D. Sardana, K. Yadav, H. Shweta, N. S. Clovis, P. Alam and S. Sen, Origin of Slow Solvation Dynamics in DNA: DAPI in Minor Groove of Dickerson-Drew DNA, Journal of Physical Chemistry B, DOI:10.1021/acs.jpcb.9b09275.

23 D. Andreatta, J. L. Pérez Lustres, S. A. Kovalenko, N. P. Ernsting, C. J. Murphy, R. S. Coleman and M. A. Berg, Power-law solvation dynamics in DNA over six decades in time, J Am Chem Soc, DOI:10.1021/ja044177v.

24 B. Y. Michel, D. Dziuba, R. Benhida, A. P. Demchenko and A. Burger, Probing of Nucleic Acid Structures, Dynamics, and Interactions With Environment-Sensitive Fluorescent Labels, 2020, preprint, DOI: 10.3389/fchem.2020.00112.

25 H. Shweta and S. Sen, Dynamics of water and ions around DNA: What is so special about them?, J Biosci, DOI:10.1007/s12038-018-9771-4.

26 S. Pal, P. K. Maiti, B. Bagchi and J. T. Hynes, Multiple time scales in solvation dynamics of DNA in aqueous solution: The role of water, counterions, and cross-correlations, Journal of Physical Chemistry B, DOI:10.1021/jp065690t.

27 K. E. Furse and S. A. Corcelli, Molecular dynamics simulations of DNA solvation dynamics, Journal of Physical Chemistry Letters, DOI:10.1021/jz100485e.

28 P. Várnai and K. Zakrzewska, DNA and its counterions: A molecular dynamics study, Nucleic Acids Res, DOI:10.1093/nar/gkh765.

29 V. P. Denisov and B. Halle, Sequence-specific binding of counterions to B-DNA, Proc Natl Acad Sci U S A, DOI:10.1073/pnas.97.2.629.

30 S. Mukherjee, S. Mondal, S. Acharya and B. Bagchi, In search of the origin of long-time power-law decay in DNA solvation dynamics, 2018, preprint, DOI: 10.48550/arxiv.1808.10471.

31 K. Yadav, D. Sardana, H. Shweta, N. S. Clovis and S. Sen, Molecular Picture of the Effect of Cosolvent Crowding on Ligand Binding and Dispersed Solvation Dynamics in G-Quadruplex DNA, Journal of Physical Chemistry B, DOI:10.1021/acs.jpcb.1c09349.

32 S. D. Verma, N. Pal, M. K. Singh and S. Sen, Sequence-Dependent Solvation Dynamics of Minor-Groove Bound Ligand Inside Duplex-DNA, Journal of Physical Chemistry B, DOI:10.1021/acs.jpcb.5b01977.

33 S. D. Verma, N. Pal, M. K. Singh and S. Sen, Probe position-dependent counterion dynamics in DNA: Comparison of time-resolved stokes shift of groove-bound to base-stacked probes in the presence of different monovalent counterions, Journal of Physical Chemistry Letters, DOI:10.1021/jz300934x.

34 A. Jacobson, W. Leupin, E. Liepinsh and G. Otting, Minor groove hydration of DNA in aqueous solution: Sequence-dependent next neighbor effect of the hydration lifetimes in d(TTAA)2 segments measured by NMR spectroscopy, Nucleic Acids Res, DOI:10.1093/nar/24.15.2911.

35 S. Mondal and S. Bandyopadhyay, Flexibility of the Binding Regions of a Protein−DNA Complex and the Structure and Ordering of Interfacial Water, J Chem Inf Model, DOI:10.1021/acs.jcim.9b00685.

36 S. K. Sinha and S. Bandyopadhyay, Dynamic properties of water around a protein-DNA complex from molecular dynamics simulations, Journal of Chemical Physics, DOI:10.1063/1.3634004.

37 R. Sarkar, R. K. Singh and S. Roy, Hierarchical Hydration Dynamics of RNA with Nano-Water-Pool at Its Core, Journal of Physical Chemistry B, DOI:10.1021/acs.jpcb.3c03553.

38 J. H. Roh, R. M. Briber, A. Damjanovic, D. Thirumalai, S. A. Woodson and A. P. Sokolov, Dynamics of tRNA at different levels of hydration, Biophys J, DOI:10.1016/j.bpj.2008.12.3895.

39 B. P. Fingerhut, The mutual interactions of RNA, counterions and water - quantifying the electrostatics at the phosphate-water interface, Chemical Communications, DOI:10.1039/d1cc05367a.

40 T. Goel, S. Kumar and S. Maiti, Thermodynamics and solvation dynamics of BIV TAR RNA-Tat peptide interaction, Mol Biosyst, DOI:10.1039/c2mb25357g.

41 X. Ye, R. A. Kumar and D. J. Patel, Molecular recognition in the bovine immunodeficiency virus Tat peptide-TAR RNA complex, Chem Biol, DOI:10.1016/1074-5521(95)90089-6.

42 V. V Pham, C. Salguero, S. N. Khan, J. L. Meagher, W. C. Brown, N. Humbert, H. de Rocquigny, J. L. Smith and V. M. D’Souza, HIV-1 Tat interactions with cellular 7SK and viral TAR RNAs identifies dual structural mimicry, Nat Commun, DOI:10.1038/s41467-018-06591-6.

43 Lindahl, Abraham, Hess and van der Spoel, GROMACS 2019.6 Manual, Zenodo, 2020, preprint, DOI: 10.5281/zenodo.3685925.

44 J. Wang, P. Cieplak and P. A. Kollman, How well does a restrained electrostatic potential (RESP) model perform in calculating conformational energies of organic and biological molecules?, J Comput Chem, 2000, 21, 1049–1074.

45 A. Pérez, I. Marchán, D. Svozil, J. Sponer, T. E. Cheatham, C. A. Laughton and M. Orozco, Refinement of the AMBER Force Field for Nucleic Acids: Improving the Description of α/γ Conformers, Biophys J, 2007, 92, 3817–3829.

46 M. Zgarbová, M. Otyepka, J. Šponer, A. Mládek, P. Banáš, T. E. I. I. I. Cheatham and P. Jurečka, Refinement of the Cornell et al. Nucleic Acids Force Field Based on Reference Quantum Chemical Calculations of Glycosidic Torsion Profiles, J Chem Theory Comput, 2011, 7, 2886–2902.

47 W. L. Jorgensen, J. Chandrasekhar, J. D. Madura, R. W. Impey and M. L. Klein, Comparison of simple potential functions for simulating liquid water, J Chem Phys, 1983, 79, 926–935.

48 W. G. Hoover, Canonical Dynamics: Equilibrium Phase-Space Distributions, Phys. Rev. A, 1985, 31, 1695.

49 S. Nosé, A unified formulation of the constant temperature molecular dynamics methods, J Chem Phys, 1984, 81, 511–519.

50 T. Darden, D. York and L. Pedersen, Particle mesh Ewald: an N · log N method for Ewald sums in large systems, J. Chem. Phys., 1993, 98, 10089.

51 B. Hess, H. Bekker, H. J. C. Berendsen and J. G. E. M. Fraaije, LINCS: A linear constraint solver for molecular simulations, J Comput Chem, 1997, 18, 1463–1472.

52 S. Mondal, S. Mukherjee and B. Bagchi, Decomposition of total solvation energy into core, side-chains and water contributions: Role of cross correlations and protein conformational fluctuations in dynamics of hydration layer, Chem Phys Lett, DOI:10.1016/j.cplett.2017.05.001.

53 L. Nilsson and B. Halle, Molecular origin of time-dependent fluorescence shifts in proteins, Proc Natl Acad Sci U S A, DOI:10.1073/pnas.0504181102.

54 M. Feig and B. Montgomery Pettitt, A molecular simulation picture of DNA hydration around A- And B-DNA, Biopolymers, DOI:10.1002/(SICI)1097-0282(1998)48:4<199::AID-BIP2>3.0.CO;2-5.

55 M. Feig and B. M. Pettitt, Sodium and chlorine ions as part of the DNA solvation shell, Biophys J, DOI:10.1016/S0006-3495(99)77023-2.

56 R. Satange and M.-H. Hou, The role of water in mediating DNA structures with epigenetic modifications, higher-order conformations and drug–DNA interactions, RSC Chem Biol, 2025, 6, 699–720.

57 A. K. Singh, C. Wen, S. Cheng and N. Q. Vinh, Long-range DNA-water interactions, Biophys J, DOI:10.1016/j.bpj.2021.10.016.

58 M. Feig and B. M. Pettitt, Modeling high-resolution hydration patterns in correlation with DNA sequence and conformation, J Mol Biol, DOI:10.1006/jmbi.1998.2486.

59 E. Frezza, D. Laage and E. Duboué-Dijon, Molecular Origin of Distinct Hydration Dynamics in Double Helical DNA and RNA Sequences, J Phys Chem Lett, 2024, 15, 4351–4358.

60 P. Auffinger and E. Westhof, RNA solvation: A molecular dynamics simulation perspective, 2000, preprint, DOI: 10.1002/1097-0282(2000)56:4<266::AID-BIP10027>3.0.CO;2-3.

61 B. Bagchi, Ed., in Water in Biological and Chemical Processes: From Structure and Dynamics to Function, Cambridge University Press, Cambridge, 2013, pp. 151–166.

62 D. Bhowmik, P. Ganesh, B. G. Sumpter and M. Goswami, Dynamical disparity between hydration shell water and RNA in a hydrated RNA system, Phys Rev E, DOI:10.1103/PhysRevE.98.062407.

63 A. Chakraborty, D. Samant, R. Sarkar, S. Sangeet, S. Prusty and S. Roy, RNA’s Dynamic Conformational Selection and Entropic Allosteric Mechanism in Controlling Cascade Protein Binding Events, J Phys Chem Lett, 2024, 15, 6115–6125.

64 D. Samant, A. Chakraborty, A. Sinha, R. Sarkar and S. Roy, Mapping Allosteric Rewiring in Viral RNA: Sequence-Encoded Control of Protein Binding Mechanisms, 2025, preprint, DOI: 10.1101/2025.11.01.685830.

65 J. Shi, J. H. Cho and W. Hwang, Heterogeneous and Allosteric Role of Surface Hydration for Protein-Ligand Binding, J Chem Theory Comput, DOI:10.1021/acs.jctc.2c00776.

66 H. Lammert, A. Wang, U. Mohanty and J. N. Onuchic, RNA as a Complex Polymer with Coupled Dynamics of Ions and Water in the Outer Solvation Sphere, Journal of Physical Chemistry B, DOI:10.1021/acs.jpcb.8b06874.

67 K. E. Furse and S. A. Corcelli, Dynamical signature of abasic damage in DNA, J Am Chem Soc, DOI:10.1021/ja109714v.

68 L. Chen and A. D. Frankel, An RNA-Binding Peptide from Bovine Immunodeficiency Virus Tat Protein Recognizes an Unusual RNA Structure, Biochemistry, DOI:10.1021/bi00175a046.

69 M. Barboric, R. Taube, N. Nekrep, K. Fujinaga and B. M. Peterlin, Binding of Tat to TAR and Recruitment of Positive Transcription Elongation Factor b Occur Independently in Bovine Immunodeficiency Virus, J Virol, DOI:10.1128/jvi.74.13.6039-6044.2000.

70 A. A. Anashkina, Protein-DNA recognition mechanisms and specificity, 2023, preprint, DOI: 10.1007/s12551-023-01137-7.

71 A. Barik and R. P. Bahadur, Hydration of protein-RNA recognition sites, Nucleic Acids Res, DOI:10.1093/nar/gku679.

