## Supplemental_Information for "Conformational and Environmental Determinants of RNA Solvation Dynamics: Roles of Intrinsic Flexibility, Allostery, and Protein Binding"

**This PDF file includes:**

**Supporting Results**

- Comparison of probability distribution of water’s residence time around TAR RNA in BIV apo to complex state.
- Power-law fitting of solvation decay computed for BIV complex state
- Computing solvation correlation function and timescale prediction at 1ps resolution and timescale prediction
- Comparison of solvation correlation function, Cs(t), decay profiles for BIV TAR-TAT complex with varying trajectories
- Solvation dynamics calculated for HIV-2 TAR and HIV-2 TAR-TAT complex and respective timescale is predicated upon fitting the plots using triexponential function
- Periodic energy fluctuation and distance fluctuation happening coherently for HIV-2 Apo
- Decomposition of solvation energy relaxation into distinct self and cross terms for the HIV-2
- Comparing global characterizations solvation environment for HIV-2 Apo vs complex

**Supporting Figures: Figure S1-S8**

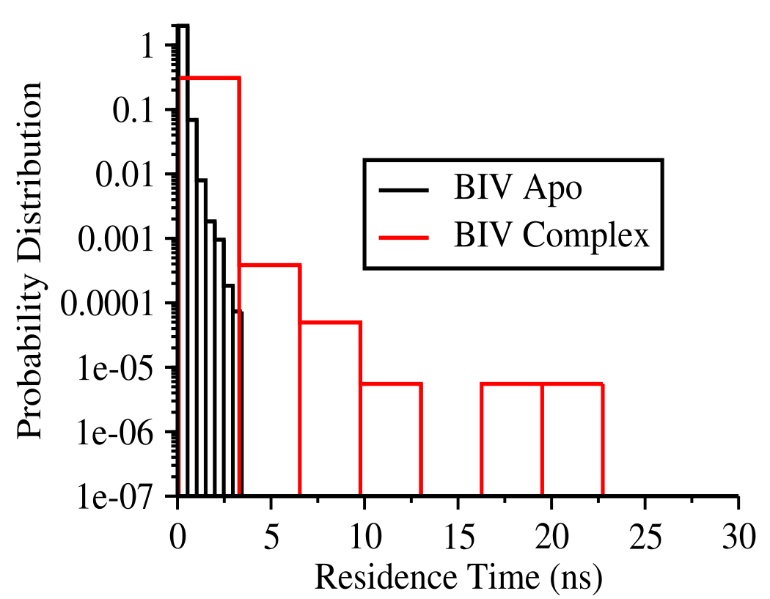
**Supporting Tables: Table S1-S5**

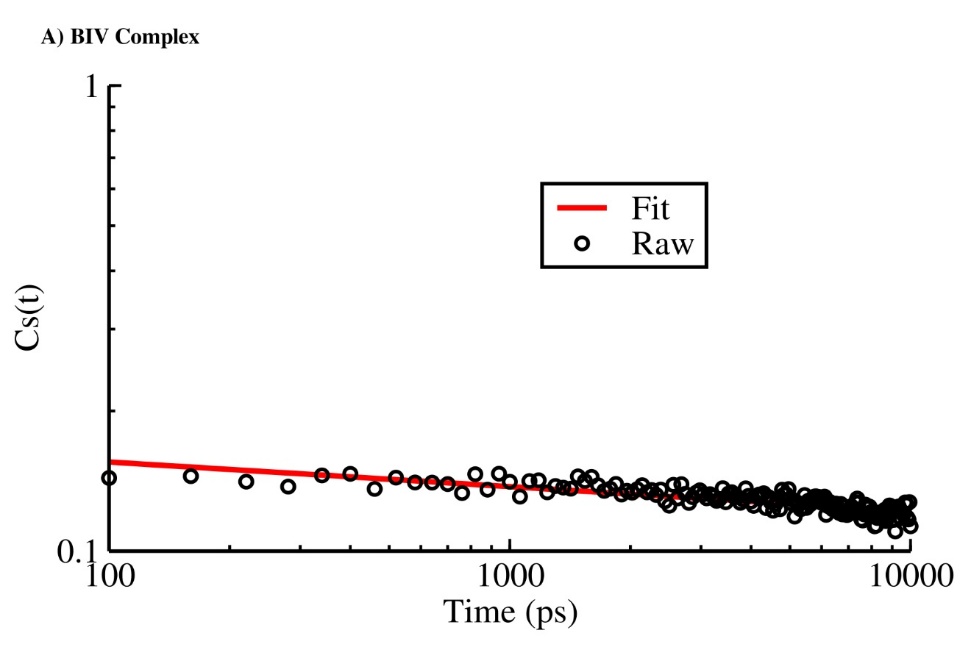
**Figure S1: Comparison of the residence time of water molecules in the first solvation shell of BIV TAR RNA in the apo state versus the TAT-bound complex.**

**Figure S2: Power-law fitting of the decay profile of the BIV TAR–TAT complex within the 100 ps to 10 ns time window (shown on a log–log scale).**

**Table S1: Fitting parameter for BIV TAR-TAT Complex (Power-Law)**

| **A0** | **A1** |
| --- | --- |
| **0.198** | **0.0529** |

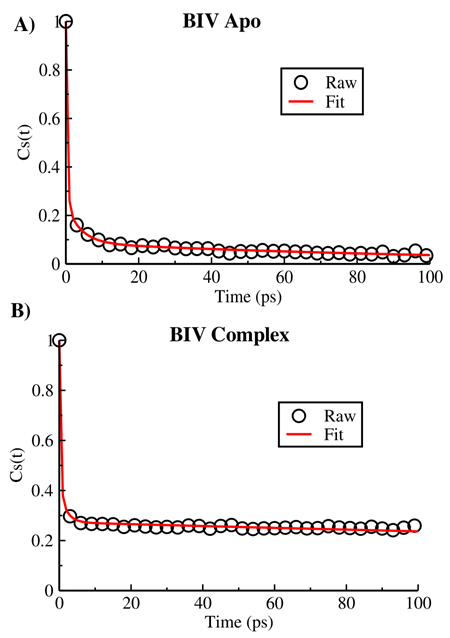

**Figure S3: Multi-exponential fitting of BIV TAR solvation dynamics at 1 ps resolution for (A) the apo state and (B) the TAT-bound complex state.**

**Table S2: Fitting parameter for BIV Apo (Tri-exponential, resolution=1ps)**

| **a1** | **a2** | **a3** | **T1** | **T2** | **T3** | **<T>** |
| --- | --- | --- | --- | --- | --- | --- |
| **0.751** | **0.162** | **0.087** | **0.36 ps** | **4.02 ps** | **114.43 ps** | **10.88 ps** |

**Table S3: Fitting parameter for BIV Complex (Tri-exponential, resolution=1ps)**

| **a1** | **a2** | **a3** | **T1** | **T2** | **T3** | **<T>** |
| --- | --- | --- | --- | --- | --- | --- |
| **0.589** | **0.138** | **0.273** | **0.32 ps** | **1.86 ps** | **700.0 ps** | **191.55 ps** |

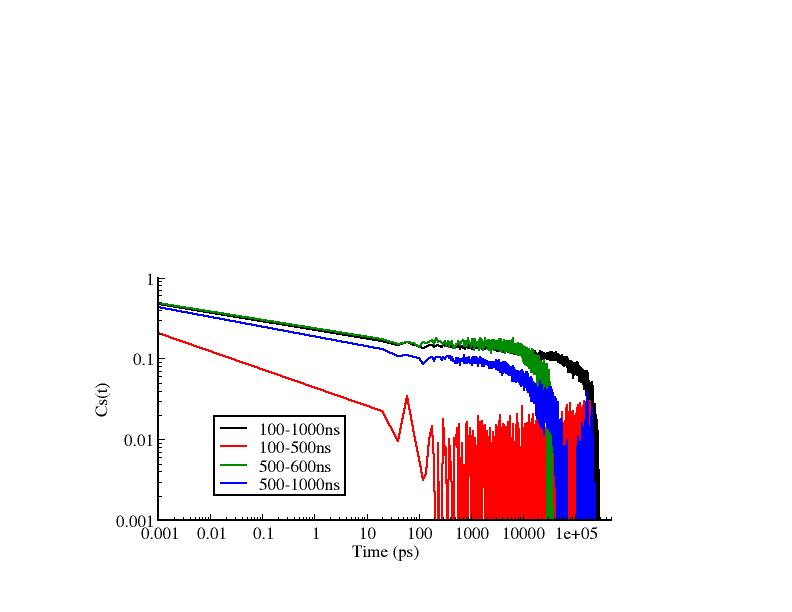

**Figure S4: Comparison of solvation time-correlation function Cs(t) decay profiles for the BIV TAR RNA in the TAT-bound complex, evaluated across trajectory segments of varying lengths (100–1000 ns, 100–500 ns, 500–600 ns, and 500–1000 ns).**

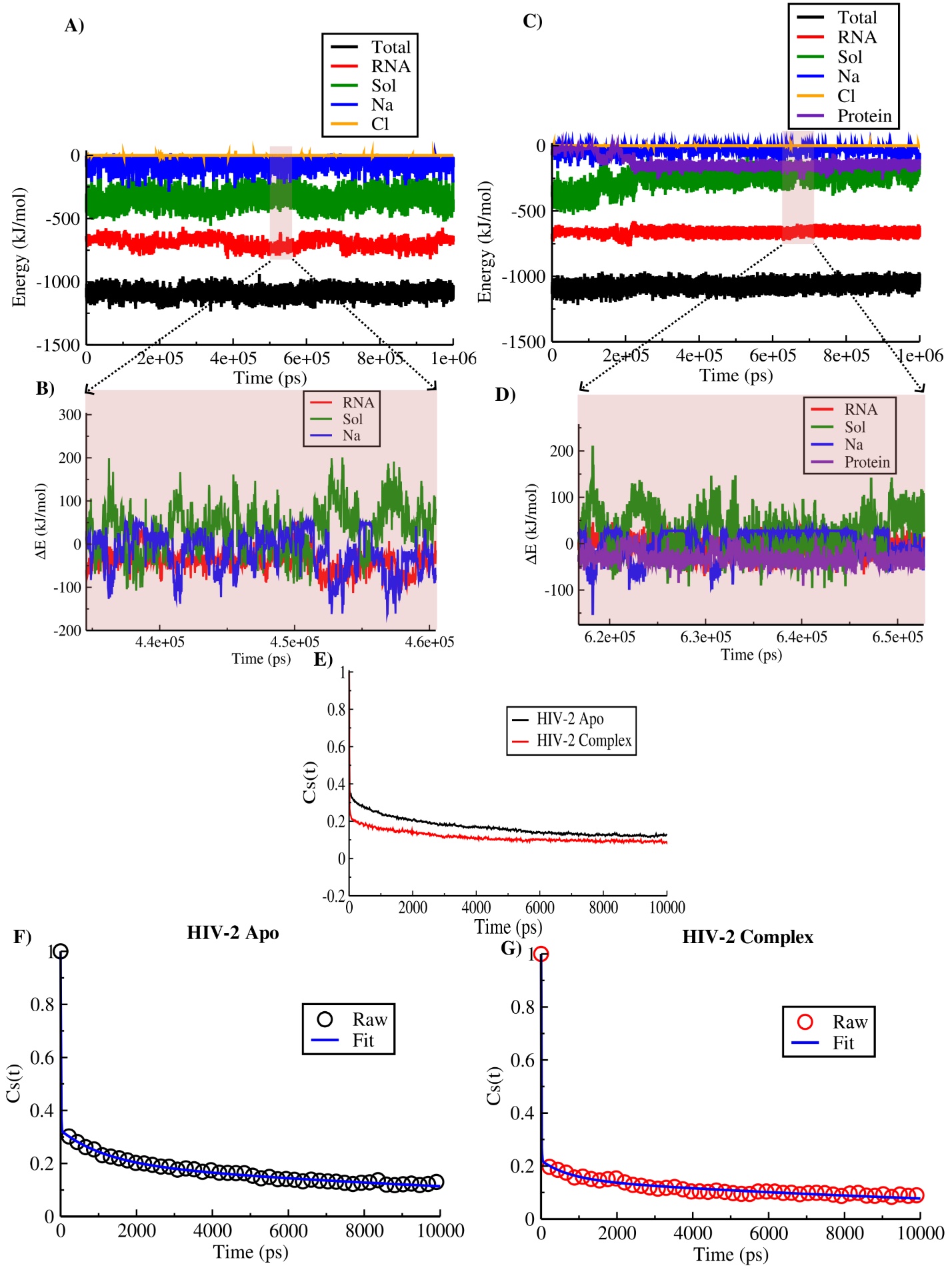

**Figure S5: Decomposition of solvation energy relaxation into distinct environmental contributions for the HIV-2 TAR RNA in its apo and TAT-bound complex states.**

(A) Probe–rest interaction energy along the trajectory for the HIV-2 apo RNA, shown as both the total contribution and its decomposition into individual components. (B) Enlarged view of panel (A), highlighting fluctuations in interaction energy (ΔE) for the RNA, solvent, and Na⁺ ion components. (C) and (D) Same analyses as in panels (A) and (B), respectively, but for the HIV-2 TAR–TAT complex. (E) Comparative decay profiles of the solvation time-correlation function for the apo and complex systems at 20 ps resolution. (F) and (G) Triexponential fits to the solvation decay profiles for the apo and complex states, respectively.

**Table S4: Fitting parameter for HIV-2 Apo (Tri-exponential, resolution=20ps)**

| **a1** | **a2** | **a3** | **T1** | **T2** | **T3** | **<T>** |
| --- | --- | --- | --- | --- | --- | --- |
| **0.675** | **0.122** | **0.203** | **7.38 ps** | **1184.66 ps** | **17377.2 ps** | **3677.08 ps** |

**Table S5: Fitting parameter for HIV-2 Complex (Tri-exponential, resolution=20ps)**

| **a1** | **a2** | **a3** | **T1** | **T2** | **T3** | **<T>** |
| --- | --- | --- | --- | --- | --- | --- |
| **0.779** | **0.07** | **0.151** | **7.32 ps** | **746.97ps** | **15000 ps** | **2322.99 ps** |

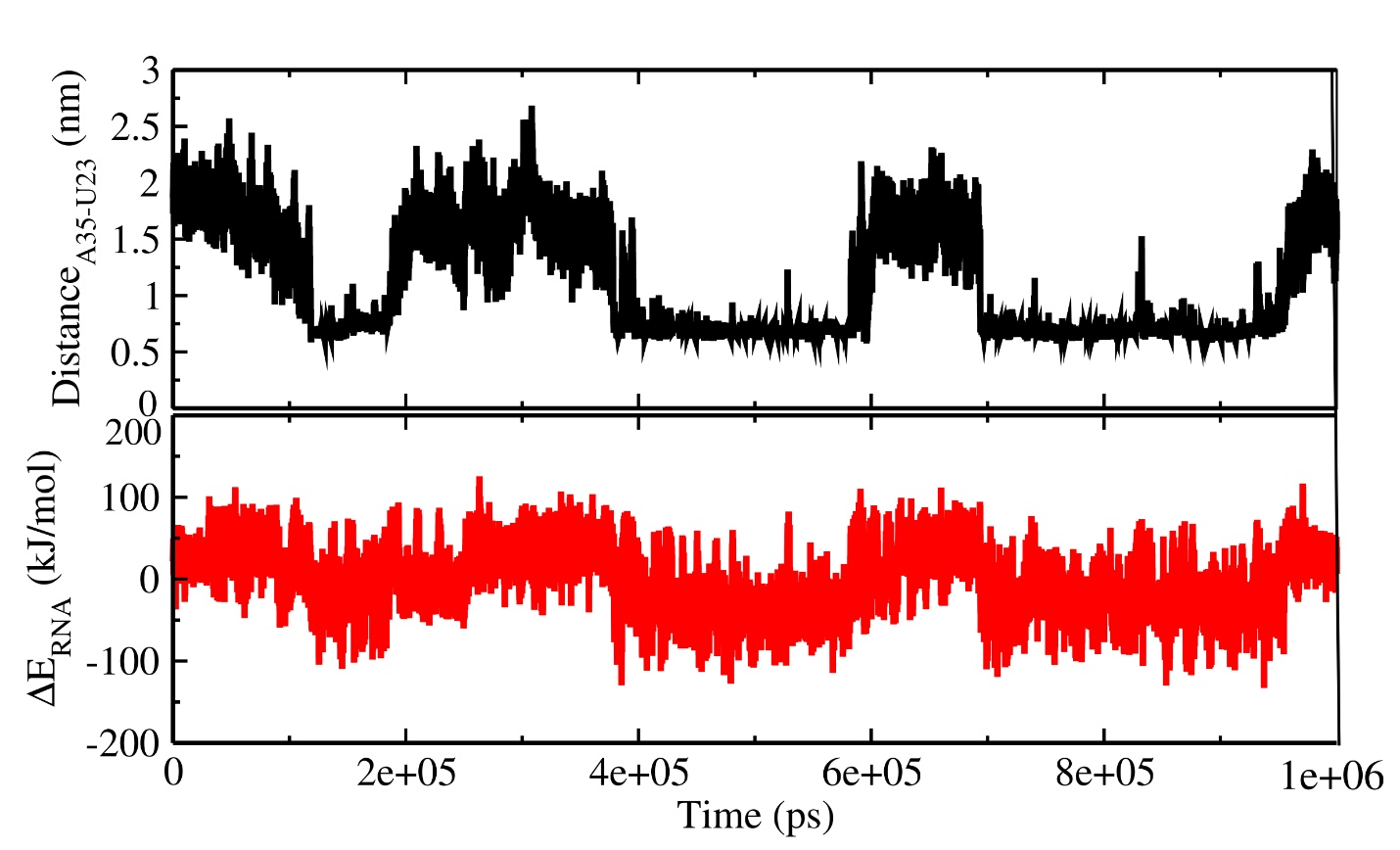

**Figure S6: Time evolution of structural energy fluctuations and their coherent periodicity with the A35 distance fluctuation, illustrating a distal conformational response sensed by the probe.**

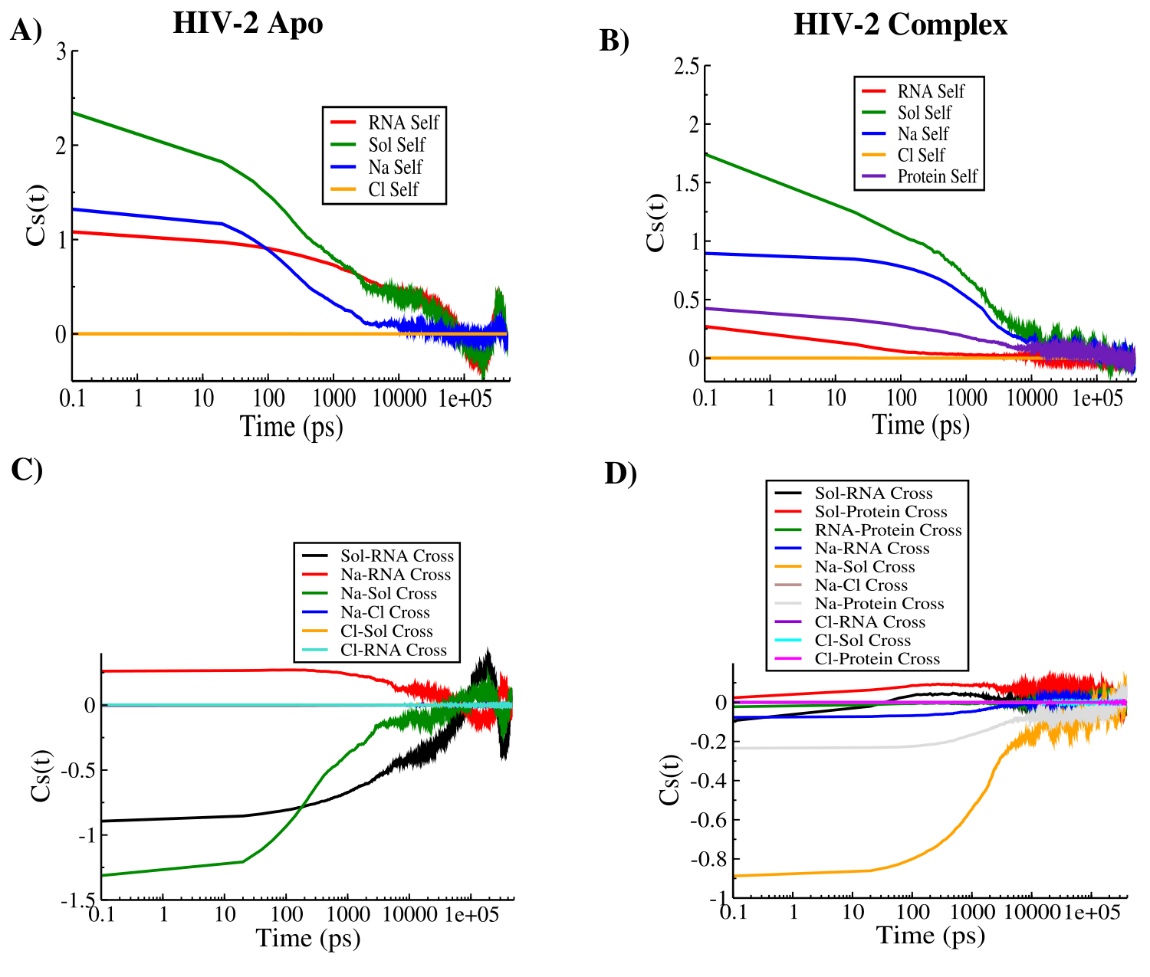

**Figure S7: Linear-response decomposition of the decay function of HIV-2 TAR RNA into self and cross terms.** (A) and (B) Self-term and cross-term contributions of the individual components governing the decay in the HIV-2 apo TAR. (C) and (D) Corresponding self-term and cross-term decompositions for the HIV-2 TAR–TAT complex.

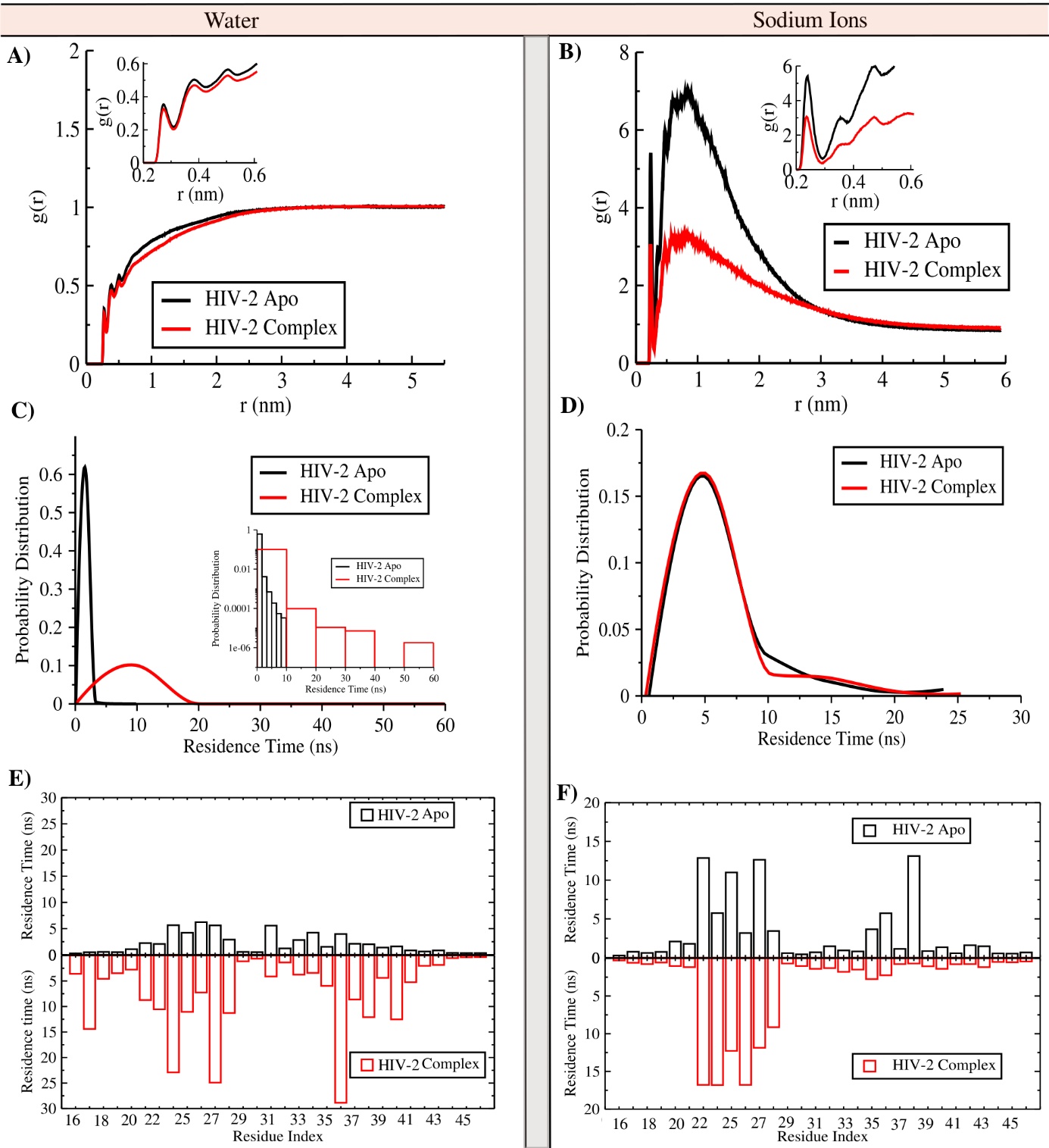

**Figure S8: Characterization of the water and ion (Na⁺) environment surrounding the RNA duplex in the apo and TAT-bound HIV-2 TAR states.** (A) and (B) Radial distribution functions of water and Na⁺ ions, respectively, around all RNA heavy atoms, shown in a comparative manner for the apo and complex forms. (C) and (D) Distributions of residence times for water and Na⁺ ions within the first hydration shell (up to 3.5 Å) of the RNA in both states. (E) and (F) Changes in the residue-wise maximum residence times of first-shell water and Na⁺ ions, respectively, upon transition from the apo state to the TAT-bound complex.
